# Identification of a chemical probe for lipid kinase phosphatidylinositol-5-phosphate 4-kinase gamma (PI5P4Kγ)

**DOI:** 10.1101/2022.09.08.507203

**Authors:** David H. Drewry, Frances M. Potjewyd, Jeffery L. Smith, Stefanie Howell, Alison D. Axtman

## Abstract

Phosphatidylinositol-5-phosphate 4-kinase gamma (PI5P4Kγ), which phosphorylates phosphatidylinositol-5-monophosphate (PI(5)P), is a human lipid kinase with intriguing roles in inflammation, T cell activation, autophagy regulation, immunity, heart failure, and several cancers. To provide a high-quality chemical tool that would enable additional characterization of this protein, we designed and evaluated a potent, selective, and cell-active inhibitor of human PI5P4Kγ. We describe the use of the PI5P4Kγ NanoBRET assay to generate structure–activity relationships (SAR), support chemical probe (**2**) design, and identify a structurally related negative control (**4**). We have characterized the binding of our chemical probe to PI5P4Kγ using orthogonal assay formats reliant on competition with an ATP-competitive reagent. Based on our results in these assays, we hypothesize that **2** binds in the ATP active site of PI5P4Kγ. Kinome-wide profiling complemented by further off-target profiling confirmed the selectivity of both our chemical probe and negative control. When a breast cancer cell line (MCF-7) was treated with compound **2**, increased mTORC1 signaling was observed, demonstrating that efficacious binding of **2** to PI5P4Kγ in cells results in activation of a negative feedback loop also reported in PI5P4Kγ knockout mice.

## 1. Introduction

Phosphatidylinositol-5-phosphate 4-kinase gamma (PI5P4Kγ) is an understudied lipid kinase belonging to the phosphatidylinositol-5-phosphate 4-kinase (PIP4K) family. PI5P4Kγ Is encoded by the gene phosphatidylinositol-5-phosphate 4-kinase type 2 gamma (*PIP4K2C/PIP5K2C*). Mammals express three PIP4K enzymes: PIP4Kα, PIP4Kβ, and PI5P4Kγ. These kinases have similar protein structure including a C-terminal kinase domain and an N-terminal dimerization domain (Rao *et al*., 1998). While PIP4Kα and PIP4Kβ share 83% identity, PI5P4Kγ only shares 60% identity with either one (Rao *et al*., 1998). Examination of the active site residues of the three isoforms revealed minor residue differences in the binding pockets and a more significant difference in the gatekeeper residues (Wortmann *et al*., 2021). Pronounced sequence divergence was observed in the P-loop that flanks the ATP site in these kinases (Wortmann *et al*., 2021).

Phosphatidylinositol-5-monophosphate (PI(5)P) is a known substrate of PIP4K family kinases and its phosphorylation produces phosphatidylinositol-4,5-bisphosphate (PI(4,5)P_2_) (Lietha, 2001; Mackey *et al*., 2014). The main function of PIP4Ks is to regulate intracellular PI(5)P and/or generate PI(4,5)P_2_ at specific locations in the cell (Sun *et al*., 2013). There is evidence for the presence of these three enzymes in the plasma membrane, Golgi, and nucleus and that each tissue has a different ratio of the proteins (Jones *et al*., 2006; Bultsma *et al*., 2010; Mackey *et al*., 2014; Wang *et al*., 2019). PI(4,5)P_2_ is a phosphoinositide that regulates membrane trafficking events such as vesicle budding and fusion, facilitates actin remodeling at the plasma membrane, and is the substrate used in generation of essential secondary messengers (Balla, 2013; Sun *et al*., 2013; Mackey *et al*., 2014; Fruman *et al*., 2017).

Despite significant catalytic site structural similarity to the protein kinase superfamily (Rao *et al*., 1998), members of the PIP4K family demonstrate drastically different catalytic rates (Clarke and Irvine, 2013). PIP4Kα is significantly more active than PIP4Kβ and PI5P4Kγ has little or no intrinsic enzymatic activity (Clarke and Irvine, 2013; Wang *et al*., 2019). These differences in activities are observed in *C. elegans, Drosophila*, and mammalian isoforms (Clarke and Irvine, 2013). It has been suggested that PI5P4Kγ serves as a chaperone for the more active isoforms, regulating their localization and/or activity (Clarke and Irvine, 2013; Shim *et al*., 2016).

The function of PI5P4Kγ is not completely characterized and that contributed to its inclusion on the understudied kinase list for the illuminating the druggable genome (IDG) project (Rodgers *et al*., 2018; Berginski *et al*., 2020). PI5P4Kγ is expressed primarily in the brain, heart, kidney, and testes (Magadum *et al*., 2021). Reports suggest roles for PI5P4Kγ in the modulation of vesicle trafficking, cell proliferation, mammalian target of rapamycin (mTOR) signaling, development and maintenance of epithelial cell functional polarity, regulation of autophagy, and eliciting analgesia (Clarke *et al*., 2008; Mackey *et al*., 2014; Sharma *et al*., 2019).

PI5P4Kγ and mammalian target of rapamycin complex 1 (mTORC1) exist in a self-regulated feedback loop (Mackey *et al*., 2014). *Pip4k2c*^*-/-*^ mice demonstrate normal growth and viability, but show hyperinflammation, increased TORC1 signaling, and enhanced T cell activation (Shim *et al*., 2016; Poli *et al*., 2021). Decreased PI5P4Kγ expression increases mTORC1 signaling, suggesting that PI5P4Kγ inhibitors could enhance immunotherapeutic strategies (Mackey *et al*., 2014; Shim *et al*., 2016). Accordingly, a human single nucleotide polymorphism (SNP) near *PIP4K2C* is associated with increased susceptibility to autoimmune diseases (Raychaudhuri *et al*., 2008). *PIP4K2C* is significantly downregulated in the hearts of cardiac hypertrophy and heart failure patients when compared to non-injured hearts (Magadum *et al*., 2021). Reduced *PIP4K2C* mRNA levels have been associated with favorable clinical outcomes of AML patients, suggesting that PI5P4Kγ inhibitors could act as anti-cancer therapeutics (Lima *et al*., 2019). Recently it has been discovered that C6-ceramide binds directly to PI5P4Kγ, preventing or altering the interaction of PI5P4Kγ with mTOR (Zhang et al., 2021). These authors also report that PI5P4Kγ promotes both metastasis and proliferation of gallbladder cancer (GBC) cells, and that high expression of this enzyme is correlated with poor prognosis for GBC patients.

Allostery has been pursued as an approach to identify selective inhibitors of PI5P4Kγ. When profiled kinome-wide at 10 µM, thienopyrimidine-based inhibitor NCT-504 (Fig. 1), discovered in a phenotypic screen, binds PI5P4Kγ with a percent of control (PoC) = 4.9 and all other kinases demonstrated PoC >40. In the same assay, NCT-504 demonstrated a *K*_D_ = 354 nM. A large shift in activity was observed when NCT-504 was analyzed for its ability to inhibit phosphorylation of PI(5)P (IC_50_ = 15.8 µM). The difference in potency observed in the enzymatic versus binding assays was ascribed to the compound binding allosterically (Al-Ramahi *et al*., 2017). NIH-12848 (Fig. 1) is an allosteric inhibitor that binds in the region on PI5P4Kγ where PI(5)P binds (Clarke *et al*., 2015). When screened in a kinome-wide panel at 10 µM, it proved to be exquisitely selective, bind PI5P4Kγ with PoC = 8.8, and bind all other kinases with PoC >47 (Liang *et al*., 2014; Clarke *et al*., 2015). This PoC value translated to a *K*_D_ = 2–3 µM in the same binding assay and an equivalent IC_50_ value in the corresponding enzymatic assays (Clarke *et al*., 2015). When human regulatory T cells were treated with NIH-12848, it phenocopied *PIP4K2C* knockdown (Poli *et al*., 2021). Another group optimized NIH-12848 to furnish compound 40 (Fig. 1), which also binds allosterically. This group solved co-crystal structures of compound 40 bound to PI5P4Kγ (PDB codes: 7QIE and 7QPN), determined a *K*_D_ = 68 nM for PI5P4Kγ, and only found PAK2 as an off-target with <50% residual activity when compound 40 was screened at 10 µM in a panel of 140 protein kinases and 15 lipid kinases (Boffey *et al*., 2022).

**Fig. 1.**
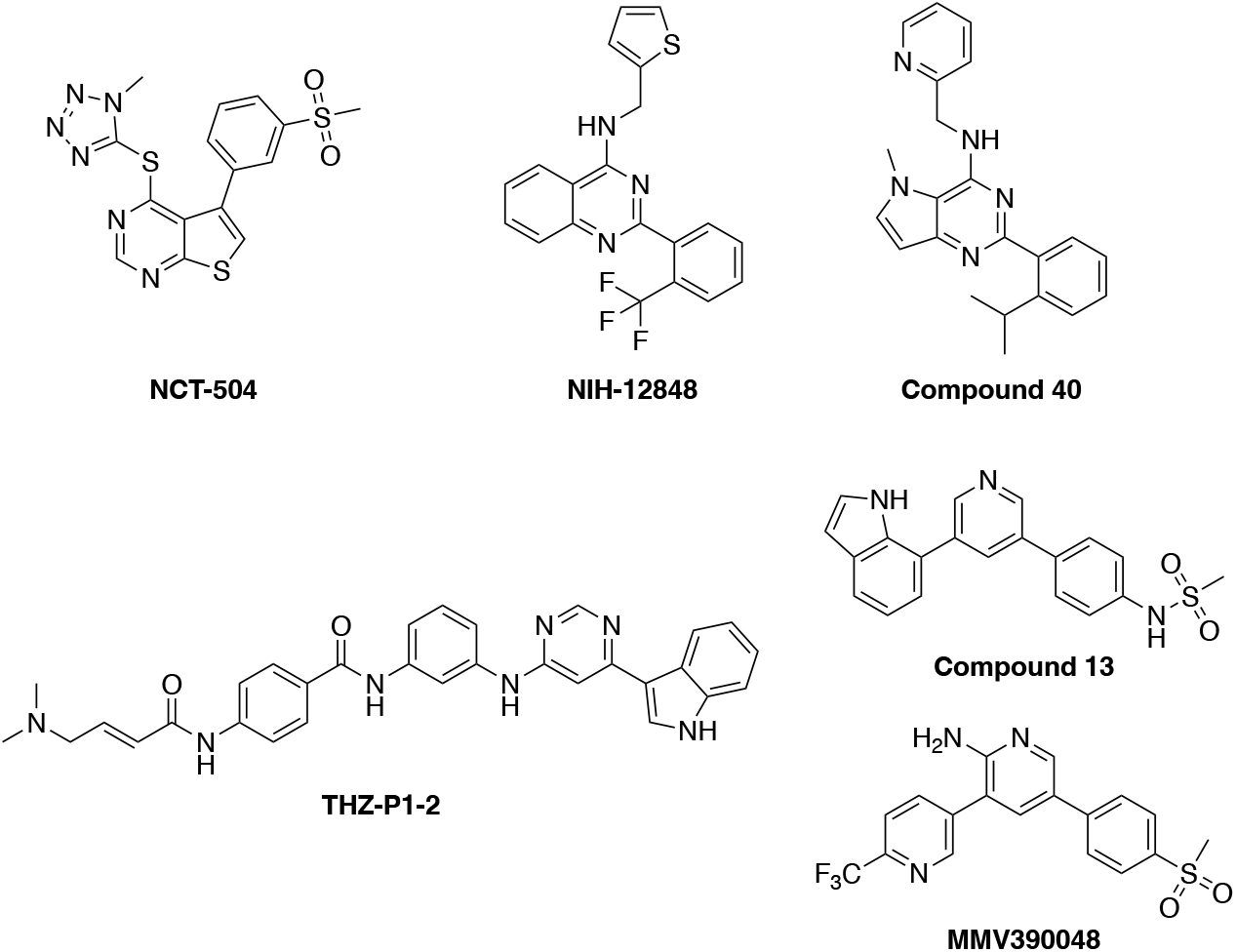
Chemical structures of allosteric PI5P4Kγ, pan-PIP4K, and P*f*PI4K inhibitors.

Pan-PIP4K inhibitors have also been reported. THZ-P1-2 (Fig. 1) was developed as a pan-PIP4K inhibitor that covalently targets cysteines on a disordered loop in PIP4Kα/β/γ, resulting in inhibition of their kinase activity at 0.2–2.8 µM. Kinome-wide selectivity of THZ-P1-2 was good (S_10_ (1 µM) = 0.02) and PI5P4Kγ demonstrated a *K*_D_ = 4.8 nM (PoC <10). The off-target kinases that demonstrated enzyme inhibition with an IC_50_ <320 nM, including ABL1, BRK, and PIKfyve, could not be pulled down in lysate (Sivakumaren *et al*., 2020). Non-covalent compound 13 (Fig. 1) has demonstrated IC_50_ = 2.0 and 9.4 µM in PIP4Kα and PIP4Kβ biochemical activity assays, respectively, and *K*_D_ = 3.4 nM (PoC = 0) when tested in a PI5P4Kγ binding assay at 1 µM (Manz *et al*., 2020). Compound 13 demonstrated excellent kinome-wide selectivity (S_35_(1 µM) = 0.02) and bound to PI5P4Kγ, RIPK2, TNNI3K, TNIK, and PIP5K1C with PoC <30 (Manz *et al*., 2020).

Attempts to make a selective ATP-competitive inhibitor of PI5P4Kγ have not been fruitful, but there are a number of inhibitory scaffolds. MMV390048 (Fig. 1) is an ATP-competitive kinase inhibitor of P*f*PI4K, the *P. falciparum* PI4K, that binds PI5P4Kγ with similar affinity (Paquet *et al*., 2017). There is clear structural similarity between compound 13 and MMV390048. PI5P4Kγ was also described as a potent off-target of LRRK2 inhibitor CZC-25146, PI3Kγ inhibitor CZC-24832, and PIP5K1C inhibitor UNC3230 (Fig. S1) (Ramsden *et al*., 2011; Bergamini *et al*., 2012; Wright *et al*., 2014). PI5P4Kγ is also an off-target of the chronic myeloid leukemia drug Gleevec (p*K*_D_ = 6.4) and urinary tract pain drug phenazopyridine (*K*_D_ = 540 nM) (Fig. S1) (Bantscheff *et al*., 2007; Preynat-Seauve *et al*., 2021). The ATP-competitive compounds commonly demonstrate binding to multiple lipid kinases (PIKfyve, PIP5K1C, PI3Kγ) in addition to PI5P4Kγ.

Here we present a potent and selective ATP-competitive PI5P4Kγ chemical probe (**2**). The tricyclic-indolyl-pyrimidinamine core is not exemplified in published inhibitors of PI5P4Kγ (Fig. 1). The PI5P4Kγ chemical probe was analyzed for binding, cellular target engagement, kinome-wide selectivity, and disruption of PI5P4Kγ-mediated signaling. A suitable negative control (**4**), which is structurally similar and demonstrates good selectivity but lacks PI5P4Kγ affinity, was also identified and evaluated. This chemical probe set can be used to expand characterization of this interesting enzyme and explore the scope of potential therapeutic roles of PI5P4Kγ.

## 2. Material and methods

### 2.1 General chemistry information

Reagents were obtained from reputable commercial vendors. Solvent was removed via rotary evaporator under reduced pressure. Thin layer chromatography was used to track reaction progress. These abbreviations are used in experimental procedures: mmol (millimoles), μmol (micromoles), mg (milligrams), mL (milliliters) and r.t. (room temperature). ^1^H NMR and/or additional microanalytical data was collected to confirm identity and assess purity of final compounds. Magnet strength for NMR spectra is included in line listings. Peak positions are listed in parts per million (ppm) and calibrated versus the shift of CD_3_OD-*d*_*4*_; coupling constants (*J* values) are reported in hertz (Hz); and multiplicities are as follows: singlet (s), doublet (d), doublet of doublets/triplets (dd/dt), doublet of doublets of triplets (ddt), triplet (t), and multiplet (m). Synthesis of final products was performed by ChemSpace LLC. These compounds were confirmed to be >95% pure by HPLC analysis upon receipt.

### 2.2 Synthesis of PI5P4Kγ chemical probe

#### 2.2.1 Suzuki coupling protocol

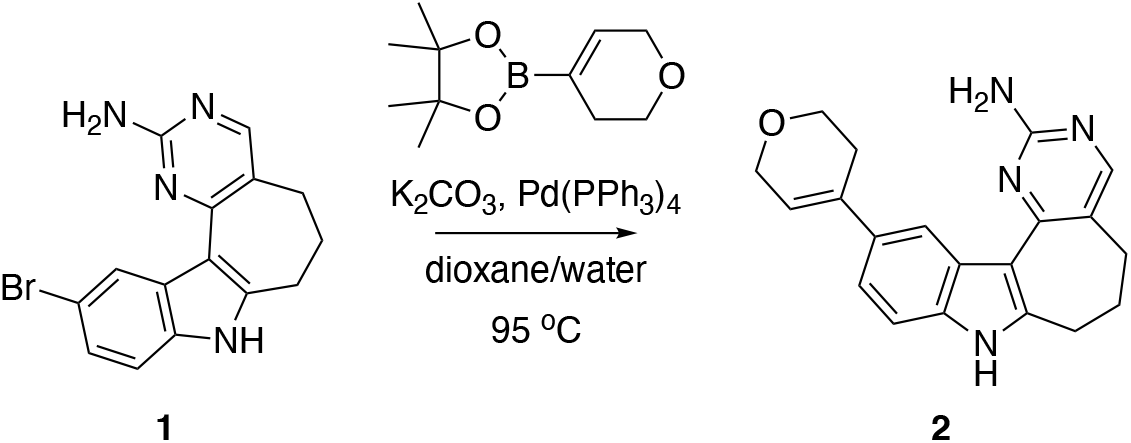

A mixture of **1** (Drewry *et al*., 2022) (192 mg, 0.58 mmol), 2-(3,6-dihydro-2*H*-pyran-4-yl)-4,4,5,5-tetramethyl-1,3,2-dioxaborolane (122 mg, 0.87 mmol), and K_2_CO_3_ (161 mg, 1.16 mmol) in a mixture of dioxane/water (2.0/0.3 mL) was degassed, followed by addition of Pd(PPh_3_)_4_ (34 mg, 29 µmol). The reaction mixture was heated to 95 °C and allowed to stir overnight. After completion of the reaction, the mixture was cooled to r.t. and diluted with ethyl acetate and water. The organic layer was washed with water and concentrated *in vacuo*. The crude residue was purified via preparative HPLC (10–100% MeOH in H_2_ O + 0.1% TFA) to afford **2** as a light yellow amorphous solid.

*11-(,6-dihydro-2H-pyran-4-yl)-5,6,7,8-tetrahydropyrimido[4’,5’:3,4]cyclohepta[1,2-b]indol-2-amine (****2***). ^1^H NMR (700 MHz, CD_3_OD-*d*_*4*_) δ 8.69 (d, *J* = 1.8 Hz, 1H), 7.88 (s, 1H), 7.32 – 7.24 (m, 2H), 6.16 – 6.14 (m, 1H), 4.33 (t, *J* = 2.7 Hz, 2H), 3.99 – 3.94 (m, 2H), 3.21 (t, *J* = 6.8 Hz, 2H), 2.73 – 2.69 (m, 2H), 2.66 (ddt, *J* = 7.7, 5.5, 2.7 Hz, 2H), 2.11 – 2.05 (m, 2H). ^13^C NMR (176 MHz, CD_3_OD-*d*_*4*_) δ 164.73, 163.06, 156.50, 145.34, 137.43, 136.75, 134.33, 129.05, 123.12, 121.02, 120.65, 120.07, 112.11, 111.13, 67.09, 65.93, 31.15, 30.60, 28.89, 26.01. HPLC purity: 100%. HRMS (ESI): *m/z* calculated for C_20_H_21_N_4_O [M + H]^+^: 333.1715. Found: 333.1709.

### 2.3 Synthesis of negative control

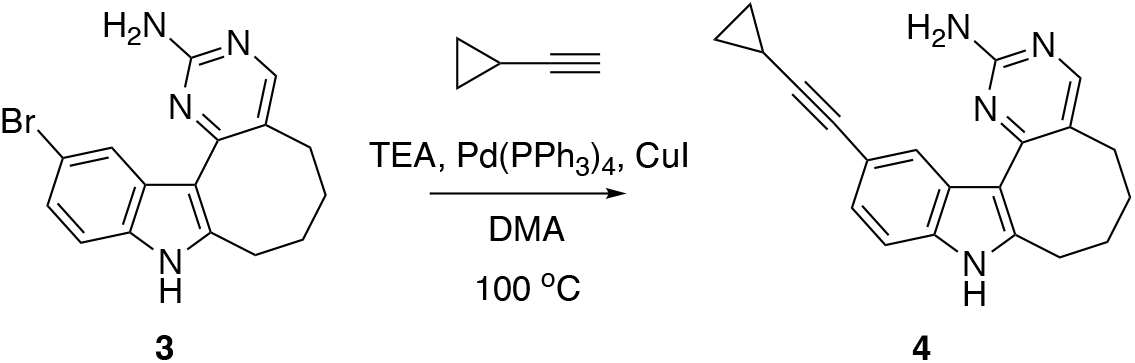

*12-(cyclopropylethynyl)-6,7,8,9-tetrahydro-5H-pyrimido[4’,5’:3,4]cycloocta[1,2-b]indol-2-amine (****4***). The analytical data for **4** matches that previously published (Drewry *et al*., 2022). ^1^H NMR (400 MHz, CD_3_OD-*d*_*4*_) δ 8.09 (s, 1H), 7.83 (s, 1H), 7.21 (dd, *J* = 8.3, 0.7 Hz, 1H), 7.10 (dd, *J* = 8.3, 1.5 Hz, 1H), 3.05 – 3.01 (m, 2H), 2.65 – 2.61 (m, 2H), 1.86 – 1.81 (m, 4H), 1.42 (dt, *J* = 8.2, 5.0 Hz, 1H), 0.87 – 0.80 (m, 2H), 0.71 – 0.66 (m, 2H).

### 2.4 Cell culture

Human embryonic kidney (HEK293) cells from ATCC were cultured in Dulbecco’s Modified Eagle’s medium (DMEM, Gibco) supplemented with 10% (v/v) fetal bovine serum (FBS, Corning). These cells were incubated at 37°C in 5% CO_2_ and passaged with trypsin (Gibco) every 72 hours to ensure that they never reached confluency.

Human breast cancer (MCF-7) cells were obtained from ATCC and cultured every two days in Minimum Essential Medium (MEM) Alpha media (Gibco) supplemented with 1 mM sodium pyruvate, 10 µg/mL human recombinant insulin, 1% non-essential amino acids, and 10% FBS. These cells were incubated at 37°C in 5% CO_2_ and passaged with trypsin (Gibco) 2–3 times per week to ensure that they never reached confluency.

### 2.5 NanoBRET assays

Constructs for NanoBRET measurements of PI5P4Kγ (PIP4K2C-NLuc), PIKfyve (PIKfyve-NLuc), MYLK4 (MYLK4-NLuc) and TYK2(JH2 domain) (TYK2(JH2 domain)-NLuc) were kindly provided by Promega. NLuc (nanoluciferase) orientations used in the corresponding assays are included in parentheses after each construct. NanoBRET assays were carried out in dose– response (12-pt curves) format as described previously (Wells *et al*., 2019; Wells *et al*., 2021). Assays according to the manufacturer’s protocol using 0.063 μM of tracer K8 for PI5P4Kγ, 0.13 μM of tracer K8 for PIKfyve, 0.13 μM of tracer K10 for MYLK4, and 0.5 μM of tracer K10 for TYK2(JH2 domain). Representative curves generated are included in Figs. S2–S4.

### 2.6 Kinome-wide selectivity analysis and confirmation of binding

The *scan*MAX assay platform offered by the Eurofins DiscoverX Corporation was used to assess the selectivity of each compound when screened at 1 µM. This platform measures the binding of a compound to 403 wild-type (WT) human as well as several mutant and non-human kinases, generating percent of control (PoC) values for every kinase evaluated (Davis *et al*., 2011). A selectivity score (S_10_ (1 µM)) is calculated using the PoC values for WT human kinases only. Kinome tree diagrams corresponding with these selectivity scores as well as tables of potently inhibited kinases are included in Figs. 2 and 3. *K*_D_ determination was carried out in dose–response format at DiscoverX for PI5P4Kγ and corresponding *K*_D_ values are incorporated into Figs. 2 and 3.

**Fig. 2.**
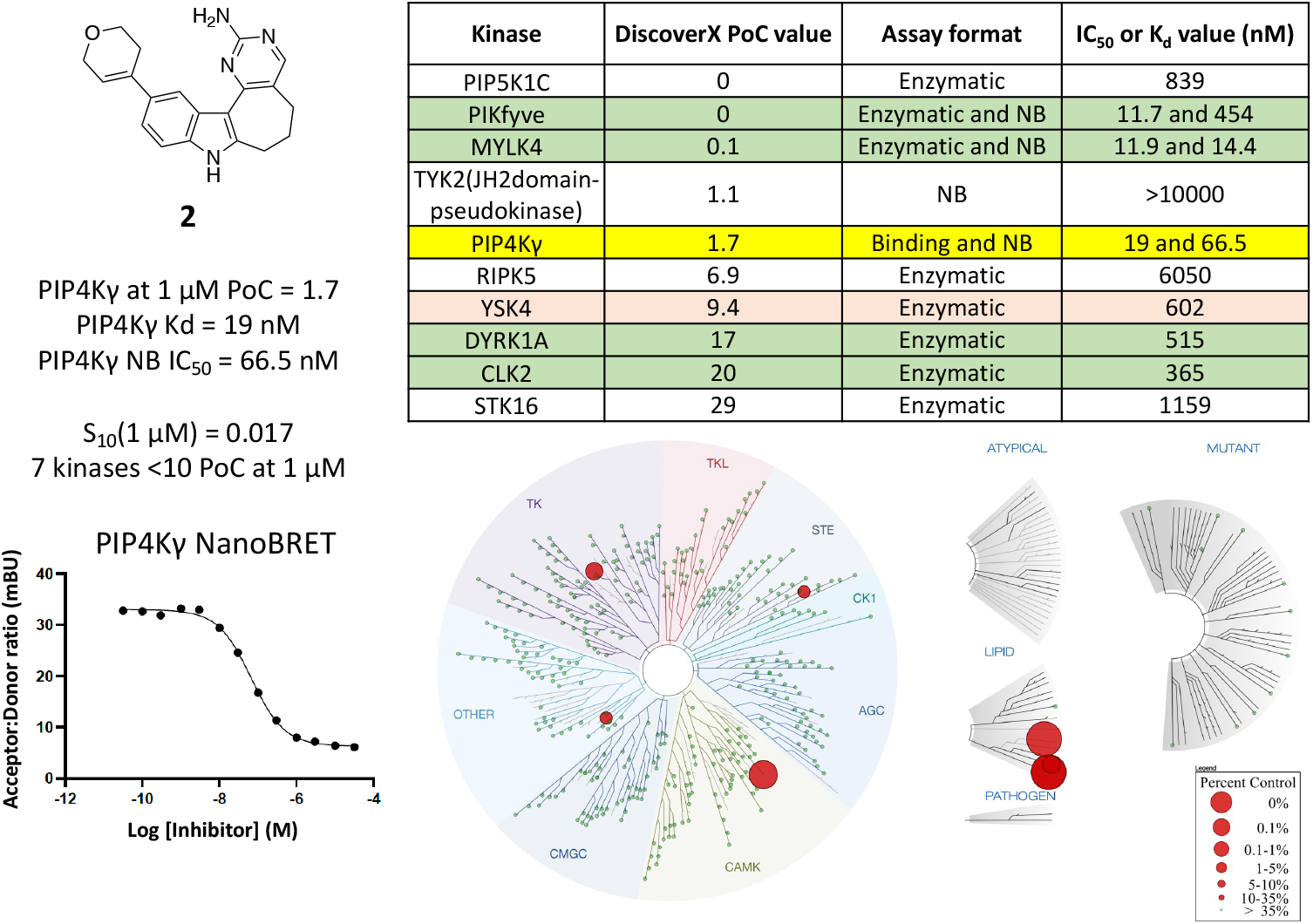
Structure, potency, and selectivity data related to PI5P4Kγ chemical probe **2**. PoC = percent of control; NB = NanoBRET. Kinome tree is representative of kinases that bind with PoC <10 when compound **2** was screened at 1 µM. Correlating assay format used to generate data in the last column of nested table is listed in the preceding column.

**Fig. 3.**
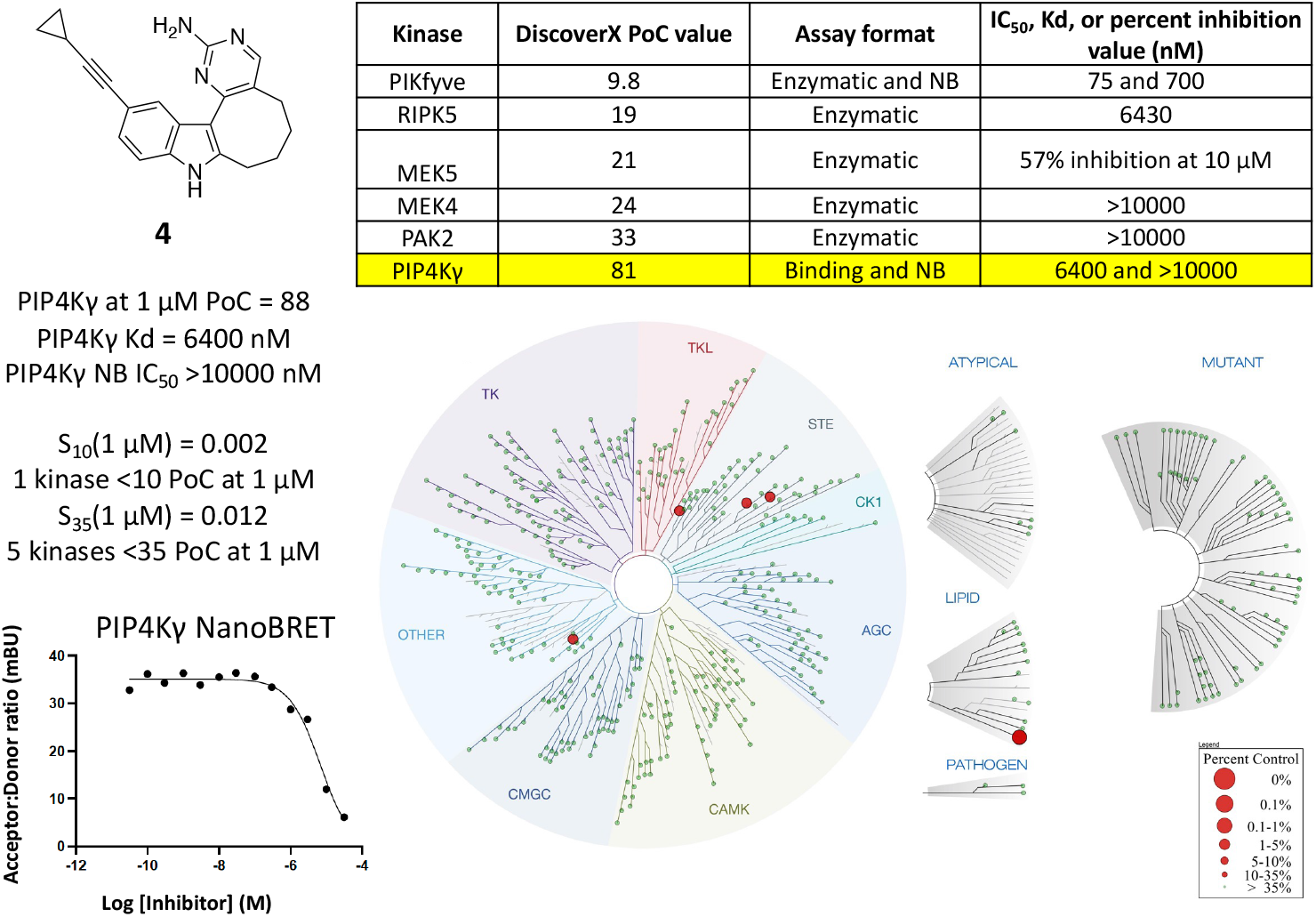
Structure, potency, and selectivity data related to PI5P4Kγ negative control **4**. PoC = percent of control; NB = NanoBRET. Kinome tree is representative of kinases that bind with PoC <35 when compound **4** was screened at 1 µM. Correlating assay format used to generate data in the last column of nested table is listed in the preceding column.

### 2.7 Orthogonal enzymatic assays

Radiometric enzymatic assays were run in dose–response (9-pt) format at Eurofins using the K_m_ value for ATP for the corresponding kinases. Specific assay details can be found on the Eurofins website: https://www.eurofinsdiscoveryservices.com. We used a previously developed ADP-Glo assay (Promega) in dose–response (10-pt curve) format in duplicate at SignalChem to evaluate the enzymatic inhibition of PIKfyve (Drewry *et al*., 2022). Radiometric HotSpot kinase assays from Reaction Biology Corp. (RBC) were carried out at the K_m_ value for ATP in dose– response (10-pt curve) format for MYLK4, RIPK5, and YSK4. Specific assay details for these HotSpot kinase assays can be viewed on the RBC website: https://www.reactionbiology.com/list-kinase-targets-us-facility. Corresponding IC_50_ values generated using these enzymatic assays are included in Fig. 2.

### 2.8 Western blot analysis

MCF-7 cells were plated at 200,000 cells per well on a 6 well plate and treated with compound **2** for 24 hours. Cells were lysed with RIPA buffer and sonicated at 50% for 10 seconds. 25 µg of protein was added to each lane on a 4-12% Tris-Glycine gel (Thermo, XP04122BOX). The gel was transferred to a PVDF membrane using the iBlot2 transfer unit. The blot was blocked in 5% milk for 1 hour at room temperature. Primary antibodies were diluted in 5% BSA; 1:5000 actin (Sigma, A2228) for 1 hour at room temperature; 1:1000 p70-S6K (p70-S6 kinase, Cell Signaling Technology, 9202S) for 4 hours at room temperature; and 1:1000 phospho-p70-S6K (Thr389) (Cell Signaling Technology, 9205S) for 4 hours at room temperature. Secondary antibodies were diluted in 5% milk and incubated for 1 hour at room temperature; 1:10,000 goat anti-mouse IRDYE 800CW (LI-COR, 926-32210); and 1:5000 goat anti-rabbit IgG HRP (Thermo, 32460). Blots were imaged on the iBright Imaging System (Thermo). Bands were analyzed using ImageJ (Fiji). Target protein bands were normalized to actin on their respective blots. Phospho-p70-S6K (Thr389) was then normalized to p70-S6K. Graphs were generated using GraphPad Prism and error bars represent standard error of the mean (SEM). Statistics were performed on 3 biological replicates using a T-test with Welch’s correction to generate p-values.

## 3. Results and discussion

### 3.1 Design of PI5P4Kγ chemical probe

Based on our published and some unpublished work, the indolyl pyrimidinamine core is a privileged lipid-targeting scaffold. We previously carried out an extensive study to establish structure–activity relationships (SAR) for this core in our development of a chemical probe targeting the lipid kinase PIKfyve (Drewry *et al*., 2022). Upon screening a small library of this scaffold at Eurofins DiscoverX in their *scan*MAX panel, we discovered that many of the compounds in that set bound to PI5P4Kγ, suggesting that we could design a cell-active PI5P4Kγ-targeting chemical probe by modifying this scaffold. To help confirm this hypothesis, several compounds were screened in a NanoBRET (NB) cellular target engagement assay to gauge their binding to PI5P4Kγ in cells (Fig. S2). Evaluation of biaryl (Table 1) and alkyne-bearing (Table 2) analogs confirmed that the binding affinity observed for PI5P4Kγ in the DiscoverX *scan*MAX panel translated to robust cellular engagement of PI5P4Kγ. Small aromatic rings were well tolerated by PI5P4Kγ, resulting in biaryl analogs with PI5P4Kγ NB IC_50_ values <260 nM. Enhanced affinity was observed for compounds bearing a six-membered ring as part of the core (**5** and **7**) when compared to their seven-membered ring congeners (**6** and **8**). Evaluation of the alkyne-bearing analogs in Table 2 demonstrated that small groups on the alkyne terminus, when attached to cores bearing six- and seven-membered rings (**9**–**13**), resulted in compounds with PI5P4Kγ NB IC_50_ values ≤200 nM. In contrast, when bulkier groups were attached to the alkyne terminus of cores bearing six- and seven-membered rings (**14** and **16**–**18**) or a core bearing an eight-membered ring (**15** and **4**), analogs with PI5P4Kγ NB IC_50_ values ≥850 nM were generated. The affinity of biaryl analogs motivated the design of a less planar analog, which was achieved by incorporating a heteroatom-bearing, non-aromatic ring of a similar size in compound **2** (Fig. 2).

**Table 1.** PI5P4Kγ binding data for biaryl analogs

| Compound | n | R | PI5P4Ky PoC<sup>[a]</sup> | PI5P4Ky NB IC<sub>50</sub> (nM) |
| --- | --- | --- | --- | --- |
| 5 | 1 | A | 0.4 | 28.7 |
| 6 | 2 | A | 15 | 76.6 |
| 7 | 1 | B | 0.3 | 167 |
| 8 | 2 | B | 5.2 | 256 |

<sup>a</sup> Percent of control (PoC) values determined at 1 μM via DiscoverX *scan*MAX profiling.

**Table 2.**
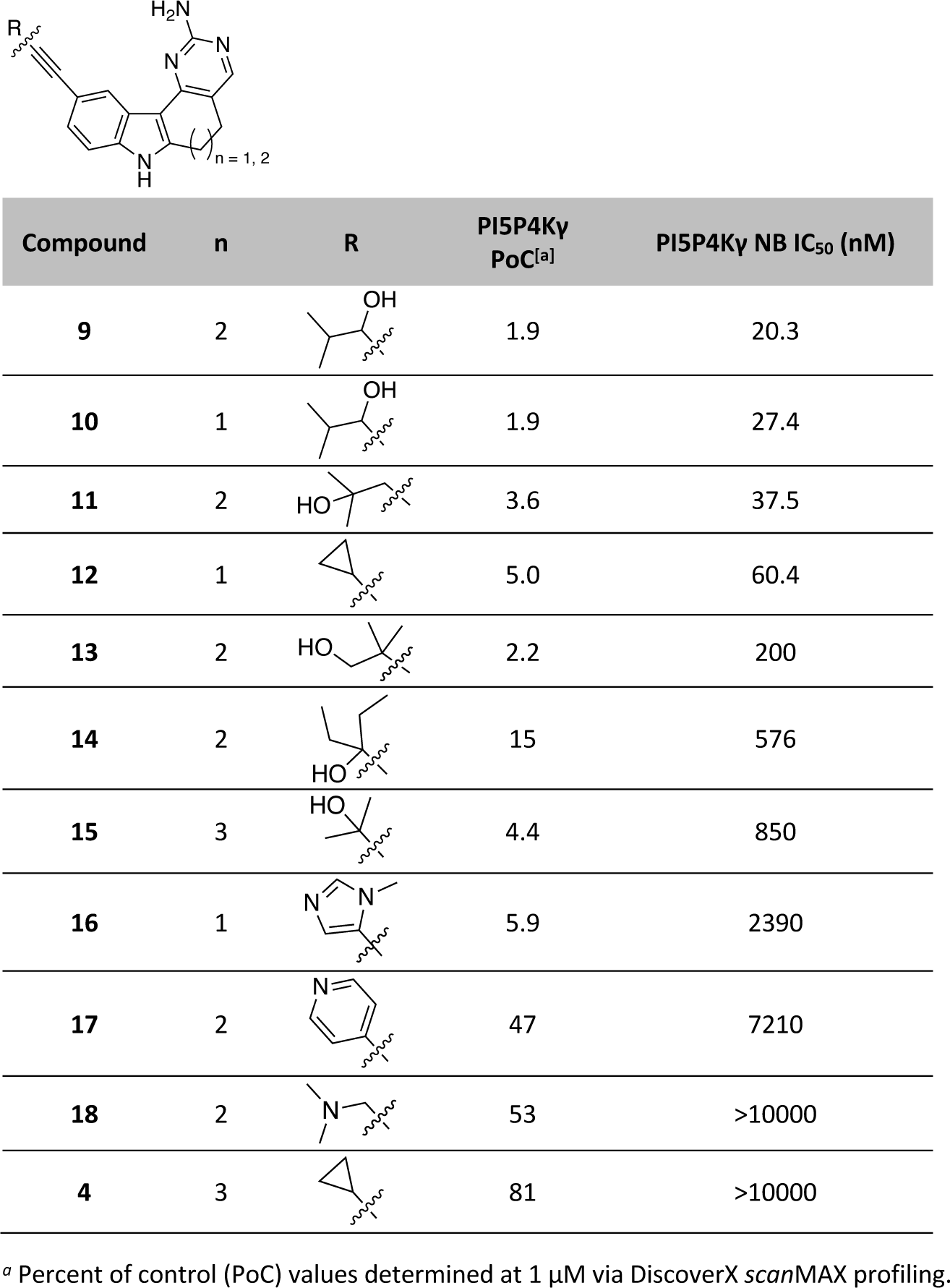
PI5P4Kγ binding data for alkyne-bearing analogs

| Compound | n | R | PI5P4Ky PoC<sup>[a]</sup> | PI5P4Ky NB IC<sub>50</sub> (nM) |
| --- | --- | --- | --- | --- |
| 9 | 2 |  | 1.9 | 20.3 |
| 10 | 1 |  | 1.9 | 27.4 |
| 11 | 2 |  | 3.6 | 37.5 |
| 12 | 1 |  | 5.0 | 60.4 |
| 13 | 2 |  | 2.2 | 200 |
| 14 | 2 |  | 15 | 576 |
| 15 | 3 |  | 4.4 | 850 |
| 16 | 1 |  | 5.9 | 2390 |
| 17 | 2 |  | 47 | 7210 |
| 18 | 2 |  | 53 | >10000 |
| 4 | 3 |  | 81 | >10000 |

<sup>a</sup> Percent of control (PoC) values determined at 1 μM via DiscoverX *scan*MAX profiling.

### 3.2 Characterization of the PI5P4Kγ chemical probe set

Using the PI5P4Kγ NB assay, compound **2** was found to potently engage with PI5P4Kγ and demonstrate an IC_50_ = 67 nM. The corresponding dose–response curve is included in Fig. 2. As a secondary confirmatory assay, the DiscoverX PI5P4Kγ binding assay was employed, and a *K*_D_ = 19 nM was observed for compound **2**. Evaluation of compound **4** in the PI5P4Kγ NB assay demonstrated that it lacks affinity for PI5P4Kγ in a cellular context (IC_50_ >10000 nM, curve in Fig. 3), consistent with the PI5P4Kγ PoC = 81 in the Eurofins DiscoverX binding assay. It was one of the lowest affinity indolyl pyrimidinamine analogs in Tables 1 and 2. Moving into a cell-free system, the DiscoverX PI5P4Kγ binding assay was used to determine a *K*_D_ = 6400 nM for compound **4**. These orthogonal assays confirmed binding of our chemical probe (**2**) to PI5P4Kγ and a lack of binding for negative control **4**. Based on the assay formats used, which both rely on displacement of an ATP-competitive molecule, we hypothesize that our chemical probe binds to the ATP site of PI5P4Kγ. This is also supported by crystal structures of this scaffold in other kinases (PDB codes: 7JXX, 4IDV, 7ZHQ, and 7ZHP) (Potjewyd *et al*., 2022). Comparison of the structures of compounds **2** and **4** confirms that they are closely structurally related, making them a suitable probe – negative control pair.

### 3.3 Evaluation of the selectivity of PI5P4Kγ chemical probe set

We evaluated the kinome-wide selectivity of our chemical probe set at 1 µM using the Eurofins DiscoverX *scan*MAX panel. Both compounds proved to have excellent selectivity. For PI5P4Kγ probe **2**, only seven human, non-mutant kinases demonstrated PoC <10. All ten human, non-mutant kinases with a PoC <35 at 1 µM were evaluated using an orthogonal enzymatic and/or binding assay. The table nested in Fig. 2 shows these kinases, ranked by their PoC value in the DiscoverX *scan*MAX panel, and the follow-up assays that were executed (Fig. S3). The data for PI5P4Kγ is highlighted in yellow. Rows corresponding to kinases that are inhibited within 30-fold of the *K*_D_ value for PI5P4Kγ are colored green, including PIKfyve, MYLK4, DYRK1A, and CLK2. YSK4, colored in beige, is the next most potently inhibited kinase (>31-fold window versus PI5P4Kγ). Interestingly, when moving into cells the binding affinity for PIKfyve (NB IC_50_ = 454 nM, Fig. S3) as an off-target is >30-fold less than that for PI5P4Kγ (NB IC_50_ = 66.5 nM), suggesting the chemical probe has better selectivity when used in a cellular context.

Negative control compound **4** is quite selective, binding to only one human, non-mutant kinase with PoC <10 and five human, non-mutant kinases with PoC <35 in this large assay panel. As was done for the chemical probe, all five kinases with a PoC <35 at 1 µM were evaluated using an orthogonal enzymatic and/or binding assay. The table nested in Fig. 3 includes these kinases, ranked by their PoC value in the DiscoverX *scan*MAX panel, and the follow-up assays that were executed. The data collected for PI5P4Kγ is included for reference and highlighted in yellow. In comparing Figs. 2 and 3, we see that PIKfyve and RIPK5 are two off-target kinases within the S_35_ (1 µM) for these two compounds, but the follow-up data indicated only very weak inhibition of RIPK5 by both the chemical probe and the negative control. With respect to PIKfyve, for compound **4**, which was recently published as compound **28** in a PIKfyve manuscript from our lab (Drewry *et al*., 2022), the enzymatic inhibition IC_50_ value (75 nM) indicated enhanced potency versus the corresponding NB inhibition IC_50_ value (700 nM, Fig. S4). Weak PIKfyve affinity is predicted when using compound **4** in cells. Considering both orthogonal validation assays, chemical probe **2** more potently binds and inhibits PIKfyve than negative control **4**. Overall, follow-up assays confirmed that our negative control does not demonstrate strong affinity for any kinase within the S_35_(1 µM) fraction and that it generally lacks affinity for the 403 WT human kinases we screened.

### 3.4 Activation of mTORC1 signaling

To start to validate that binding of our PI5P4Kγ chemical probe could impact relevant signaling pathways we looked to the literature. Reports converged on the idea that PI5P4Kγ has limited to no capacity to phosphorylate PI(5)P (Poli *et al*., 2019). In support of its very low catalytic activity, a published PI5P4Kγ enzyme assay used a final enzyme concentration of around 2.3 μM (Clarke *et al*., 2015). Other groups have been unable to determine the PI5P4Kγ activity for putative inhibitors. Finally, mutation of the PI5P4Kγ catalytic site to the corresponding PIP4Kα G-loop sequence has been an artificial approach used to increase the kinase functional activity and enable the development of an inhibition assay with a useable window (Clarke and Irvine, 2013; Boffey *et al*., 2022).

Commercial enzymatic assays for PI5P4Kγ are not available due to full-length constructs of PI5P4Kγ not being active. This was confirmed in our attempts to use 25–50 ng/reaction of the full-length protein (PIP4K2C, SignalChem P76-102CG) to phosphorylate phosphatidylinositol 5-phosphate (PI(5)P) in phosphatidylserine (PS) (PI(5)P:PS, SignalChem P428-59) in the presence of 25 µM ATP. Another vendor that had previously used the full-length PI5P4Kγ protein from SignalChem confirmed that activity was suboptimal and high concentration of enzyme was required to get enough signal, resulting an assay that was not robust enough to include in their standard services. These concepts precluded collection of any PI5P4Kγ enzyme inhibition data for our chemical probe.

Based on the idea that PI5P4Kγ serves as a chaperone for and needs to co-exist to enhance activity of other more active PIP4K isoforms, we hypothesized that we should be able to observe activity in a cellular context given the proper readout (Clarke and Irvine, 2013; Shim *et al*., 2016). With this approach in mind, we noted that several groups had demonstrated that PI5P4Kγ knockout (KO) impacts the activation of p70-S6 kinase (p70-S6K). This is not surprising given the known feedback loop that exists between mTOR and PI5P4Kγ and that p70-S6K has been described as a mTORC1 pathway marker (Mackey *et al*., 2014; Magadum *et al*., 2021). Western blot analyses from different tissues from WT versus PI5P4Kγ KO (*Pip4k2c*^*-/-*^) mice demonstrated differences in phosphorylation of p70-S6K. Specifically, phosphorylation of p70-S6K on threonine 389 (Thr389), a site modified by mTORC1, was increased in kidney, liver, brain, and muscle tissues as well as the spleen from PI5P4Kγ KO mice compared to WT mice (Shim *et al*., 2016). A similar result was observed when both phosphorylated and non-phosphorylated p70-S6K were examined 21 days after transverse aortic constriction (TAC) in PI5P4Kγ KO (*Pip4k2c*^*-/-*^) versus WT mice. Phosphorylated p70-S6K was significantly higher in PI5P4Kγ KO mice when compared to WT mice and both cohorts had elevated phosho-p70-S6K when compared to animals not subjected to TAC injury. The authors suggest that PI5P4Kγ negatively regulates the mTORC1 pathway in the heart following TAC (Magadum *et al*., 2021).

To determine whether our chemical probe impacts a signaling pathway that responds to changes in PI5P4Kγ expression, we analyzed whether compound **2** increases phosphorylation of p70-S6K. For our experiments we selected MCF-7 cells based on their high mTORC1 and modest PIP4K2C mRNA expression. After 48 hours exposure of MCF-7 cells to compound **2**, we observed a dose-responsive increase in phospho-p70-S6K and no change in expression of p70-S6K (Fig. 4). A significant 60% increase in phospho-p70-S6K relative to p70-S6K expression was observed in response to the 5 µM dose. This is consistent with the response observed in *Pip4k2c*^*-/-*^ mice. We conclude that binding of our chemical probe to PI5P4Kγ leads to a functional response, resulting in increased phosphorylation of p70-S6K and activation of the mTORC1 pathway.

**Fig. 4.**
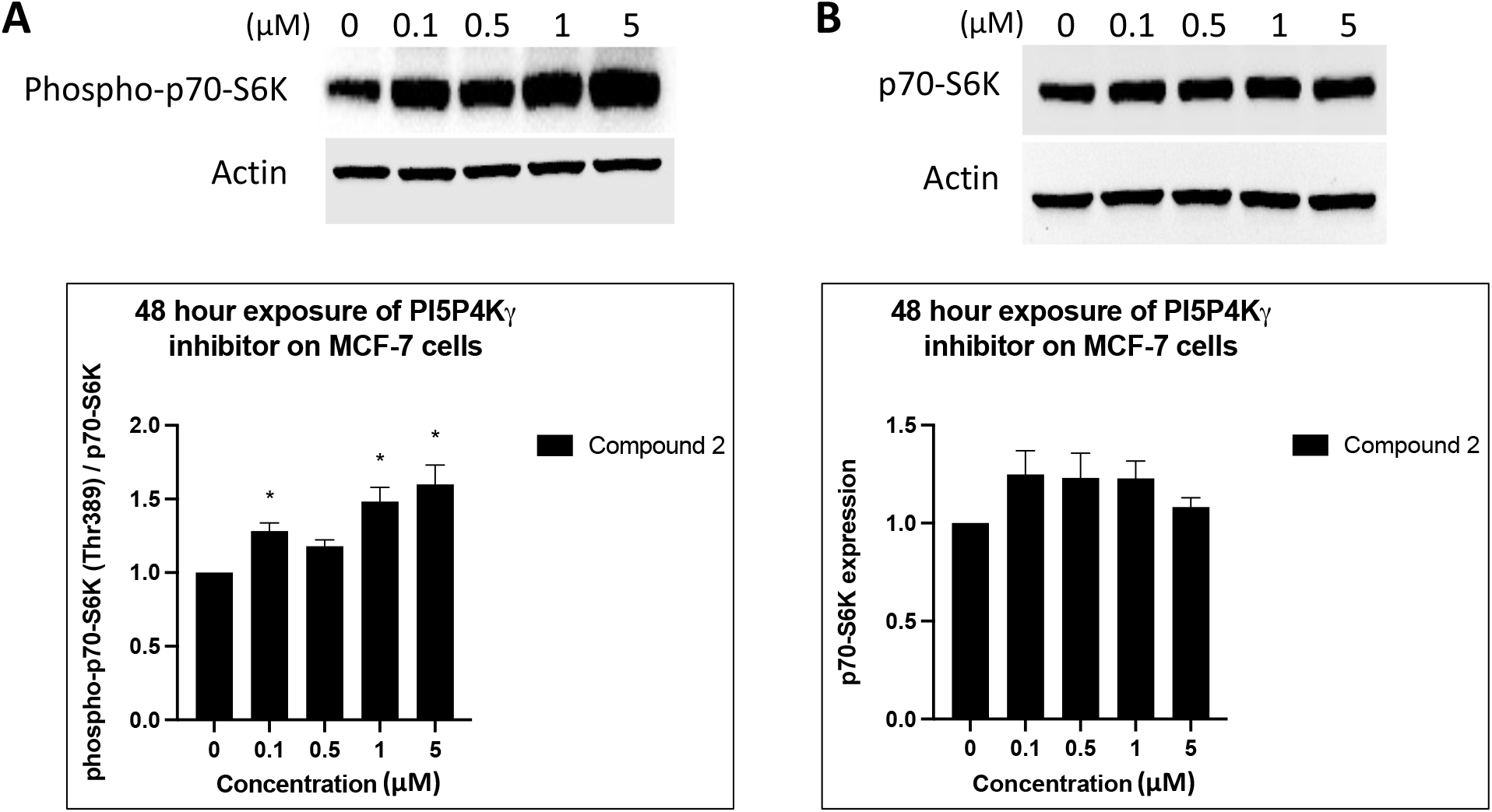
Western blot analyses of MCF-7 cells after 48 h treatment with compound **2**. (A) Representative blot and quantification of phospho-p70-S6K (Thr389, normalized to p70-S6K), n = 3. *p-values: 0.1 µM = 0.0349, 1 µM = 0.0383, and 5 µM = 0.0461. (B) representative blot and quantification of p70-S6K, n = 3. None of the doses were statistically significant compared to control. Error bars represent standard error of the mean (SEM). P-values were generated using a parametric unpaired t-test with Welch’s correction comparing each treatment condition to the untreated control.

## 4. Summary & conclusion

Indolyl pyrimidinamines **2** and **4** can be used as a chemical probe set for interrogating the function of PI5P4Kγ in cells. Compound **2** meets Structural Genomics Consortium (SGC) chemical probe criteria based upon its biochemical and cellular affinity for PI5P4Kγ : *K*_D_ = 19 nM in the PI5P4Kγ assay at DiscoverX and IC_50_ = 66.5 nM in PI5P4Kγ NB assay (Krahn *et al*., 2020). The kinome-wide selectivity of compound **2** is very good (S_10_(1 µM) = 0.017) and only four additional kinases bind within 30-fold of the PI5P4Kγ *K*_D_ value: MYLK4, PIKfyve, DYRK1A, and CLK2. Even better selectivity is observed when the chemical probe is used in cells, as PIKfyve affinity is reduced. Although a 19–27-fold selectivity window exists between PI5P4Kγ and CLK2/DYRK1A affinity, several potent and selective chemical probes targeting CLK2, with or without DYRK1A inhibition, have been described and are commercially available that can be used in tandem to rule out a phenotype due to CLK2 and/or DYRK1A inhibition: T3-CLK, MU1210, and SGC-CLK-1 (Funnell *et al*., 2017; Němec *et al*., 2019). Similarly, a PIKfyve chemical probe candidate was recently described that also has some MYLK4 inhibitory activity but lacks PI5P4Kγ affinity (Drewry *et al*., 2022). Compound **4** is a suitable negative control to be distributed and utilized alongside the PI5P4Kγ chemical probe due to its >100× lower PI5P4Kγ affinity in cell-free and intact cell binding assays. This negative control also exhibits excellent kinome-wide selectivity: S_10_(1 µM) = 0.002; S_35_(1 µM) = 0.012. Finally, chemical probe **2** was shown to phenocopy the activation of mTORC1 signaling observed in response to PI5P4Kγ KO, confirming that treatment of cells with this compound inhibits PI5P4Kγ function. We provide an ATP-competitive inhibitor of PI5P4Kγ that is active in cells and can be used to further characterize PI5P4Kγ biology.

## Supporting information

Combined Supplemental Data

## Abbreviations

*P. falciparum*: *plasmodium falciparum*
CD_3_ OD-*d*_*4*_: deuterated methanol
RIPA: radioimmunoprecipitation buffer
PVDF: polyvinylidene difluoride
BSA: bovine serum albumin
HRP: horseradish peroxidase
PDB: protein data bank
AML: acute myeloid leukemia

## Acknowledgements

NanoBRET constructs for PI5P4Kγ, PIKfyve, MYLK4, and TYK2 were provided in-kind by Promega. We thank SignalChem Biotech Inc. for helping to develop a PIKfyve enzymatic assay (Drewry *et al*., 2022) and providing PI5P4Kγ protein for enzymatic studies. The TREE*spot* kinase interaction mapping software was employed in preparation of the kinome trees in Figs 2 and 3: http://treespot.discoverx.com. Our graphical abstract was created using Biorender.com. We thank ChemSpace for synthetic support.

The Structural Genomics Consortium is a registered charity (number 1097737) that receives funds from AbbVie, Bayer Pharma AG, Boehringer Ingelheim, Canada Foundation for Innovation, Eshelman Institute for Innovation, Genome Canada, Genentech, Innovative Medicines Initiative (EU/EFPIA), Janssen, Merck KGaA Darmstadt Germany, MSD, Novartis Pharma AG, Ontario Ministry of Economic Development and Innovation, Pfizer, São Paulo Research Foundation-FAPESP, Takeda, and Wellcome. Research reported in this publication was supported in part by the NC Biotechnology Center Institutional Support Grant 2018-IDG-1030 and NIH U24DK116204.

## Appendix A. Supplementary data

Supplementary data to this article can be found online at https://doi.org/TBD.

## References

Al-Ramahi, I., Giridharan, S.S.P., Chen, Y.C., Patnaik, S., Safren, N., Hasegawa, J., de Haro, M., Wagner Gee, A.K., Titus, S.A., Jeong, H., Clarke, J., Krainc, D., Zheng, W., Irvine, R.F., Barmada, S., Ferrer, M., Southall, N., Weisman, L.S., Botas, J., Marugan, J.J., 2017. Inhibition of PIP4kγ ameliorates the pathological effects of mutant huntingtin protein. Elife, 6, e29123.

Balla, T., 2013. Phosphoinositides: Tiny lipids with giant impact on cell regulation. Physiol Rev, 93, 1019–1137.

Bantscheff, M., Eberhard, D., Abraham, Y., Bastuck, S., Boesche, M., Hobson, S., Mathieson, T., Perrin, J., Raida, M., Rau, C., Reader, V., Sweetman, G., Bauer, A., Bouwmeester, T., Hopf, C., Kruse, U., Neubauer, G., Ramsden, N., Rick, J., Kuster, B., Drewes, G., 2007. Quantitative chemical proteomics reveals mechanisms of action of clinical ABL kinase inhibitors. Nat Biotechnol, 25, 1035–1044.

Bergamini, G., Bell, K., Shimamura, S., Werner, T., Cansfield, A., Müller, K., Perrin, J., Rau, C., Ellard, K., Hopf, C., Doce, C., Leggate, D., Mangano, R., Mathieson, T., O’Mahony, A., Plavec, I., Rharbaoui, F., Reinhard, F., Savitski, M.M., Ramsden, N., Hirsch, E., Drewes, G., Rausch, O., Bantscheff, M., Neubauer, G., 2012. A selective inhibitor reveals PI3Kγ dependence of T(H)17 cell differentiation. Nat Chem BIol, 8, 576–582.

Berginski, M.E., Moret, N., Liu, C., Goldfarb, D., Sorger, P.K., Gomez, S.M., 2020. The dark kinase knowledgebase: An online compendium of knowledge and experimental results of understudied kinases. Nucleic Acids Res, 49, D529–D535.

Boffey, H.K., Rooney, T.P.C., Willems, H.M.G., Edwards, S., Green, C., Howard, T., Ogg, D., Romero, T., Scott, D.E., Winpenny, D., Duce, J., Skidmore, J., Clarke, J.H., Andrews, S.P., 2022. Development of selective phosphatidylinositol 5-phosphate 4-kinase γ inhibitors with a non-ATP-competitive, allosteric binding mode. J Med Chem, 65, 3359–3370.

Bultsma, Y., Keune, W.-J., Divecha, N., 2010. PIP4Kβ interacts with and modulates nuclear localization of the high-activity PtdIns5P-4-kinase isoform PIP4Kα. Biochem J, 430, 223–235.

Clarke, J.H., Emson, P.C., Irvine, R.F., 2008. Localization of phosphatidylinositol phosphate kinase IIgamma in kidney to a membrane trafficking compartment within specialized cells of the nephron. Am J Physiol Renal Physiol, 295, F1422–1430.

Clarke, J.H., Giudici, M.L., Burke, J.E., Williams, R.L., Maloney, D.J., Marugan, J., Irvine, R.F., 2015. The function of phosphatidylinositol 5-phosphate 4-kinase γ (PI5P4Kγ) explored using a specific inhibitor that targets the PI5P-binding site. Biochem J, 466, 359–367.

Clarke, J.H., Irvine, R.F., 2013. Evolutionarily conserved structural changes in phosphatidylinositol 5-phosphate 4-kinase (PI5P4K) isoforms are responsible for differences in enzyme activity and localization. Biochem J, 454, 49–57.

Davis, M.I., Hunt, J.P., Herrgard, S., Ciceri, P., Wodicka, L.M., Pallares, G., Hocker, M., Treiber, D.K., Zarrinkar, P.P., 2011. Comprehensive analysis of kinase inhibitor selectivity. Nat Biotechnol, 29, 1046–1051.

Drewry, D.H., Potjewyd, F.M., Smith, J.L., Dickmander, R.J., Bayati, A., Howell, S., Taft-Benz, S.A., Min, S.M., Hossain, M.A., Heise, M.T., McPherson, P.S., Moorman, N.J., Axtman, A.D., 2022. Identification and utilization of a chemical probe to interrogate the roles of PIKfyve in the lifecycle of β-coronaviruses. J Med Chem, ASAP.

Fruman, D.A., Chiu, H., Hopkins, B.D., Bagrodia, S., Cantley, L.C., Abraham, R.T., 2017. The PI3K pathway in human disease. Cell, 170, 605–635.

Funnell, T., Tasaki, S., Oloumi, A., Araki, S., Kong, E., Yap, D., Nakayama, Y., Hughes, C.S., Cheng, S.W.G., Tozaki, H., Iwatani, M., Sasaki, S., Ohashi, T., Miyazaki, T., Morishita, N., Morishita, D., Ogasawara-Shimizu, M., Ohori, M., Nakao, S., Karashima, M., Sano, M., Murai, A., Nomura, T., Uchiyama, N., Kawamoto, T., Hara, R., Nakanishi, O., Shumansky, K., Rosner, J., Wan, A., McKinney, S., Morin, G.B., Nakanishi, A., Shah, S., Toyoshiba, H., Aparicio, S., 2017. CLK-dependent exon recognition and conjoined gene formation revealed with a novel small molecule inhibitor. Nat Commun, 8, 7.

Jones, D.R., Bultsma, Y., Keune, W.-J., Halstead, J.R., Elouarrat, D., Mohammed, S., Heck, A.J., D’Santos, C.S., Divecha, N., 2006. Nuclear PtdIns5P as a transducer of stress signaling: An in vivo role for PIP4Kbeta. Mol Cell, 23, 685–695.

Krahn, A.I., Wells, C., Drewry, D.H., Beitel, L.K., Durcan, T.M., Axtman, A.D., 2020. Defining the neural kinome: Strategies and opportunities for small molecule drug discovery to target neurodegnerative diseases. ACS Chem Neurosci, 11, 1871–1886.

Liang, Q., Dexheimer, T.S., Zhang, P., Rosenthal, A.S., Villamil, M.A., You, C., Zhang, Q., Chen, J., Ott, C.A., Sun, H., Luci, D.K., Yuan, B., Simeonov, A., Jadhav, A., Xiao, H., Wang, Y., Maloney, D.J., Zhuang, Z., 2014. A selective USP1-UAF1 inhibitor links deubiquitination to DNA damage responses. Nat Chem Biol, 10, 298–304.

Lietha, D., 2001. Phosphoinositides–the seven species: Conversion and cellular roles. In: Encyclopedia of life sciences (ELS), John Wiley & Sons, Ltd, Chichester, 1–11.

Lima, K., Coelho-Silva, J.L., Kinker, G.S., Pereira-Martins, D.A., Traina, F., Fernandes, P.A.C.M., Markus, R.P., Lucena-Araujo, A.R., Machado-Neto, J.A., 2019. PIP4K2A and PIP4K2C transcript levels are associated with cytogenetic risk and survival outcomes in acute myeloid leukemia. Cancer Genet, 233–234, 56–66.

Mackey, A.M., Sarkes, D.A., Bettencourt, I., Asara, J.M., Rameh, L.E., 2014. PIP4kγ is a substrate for mTORC1 that maintains basal mTORC1 signaling during starvation. Sci Signal, 7, ra104–ra104.

Magadum, A., Singh, N., Kurian, A.A., Sharkar, M.T.K., Sultana, N., Chepurko, E., Kaur, K., Żak, M.M., Hadas, Y., Lebeche, D., Sahoo, S., Hajjar, R., Zangi, L., 2021. Therapeutic delivery of Pip4k2c-modified mRNA attenuates cardiac hypertrophy and fibrosis in the failing heart. Adv Sci, 8, 2004661.

Manz, T.D., Sivakumaren, S.C., Ferguson, F.M., Zhang, T., Yasgar, A., Seo, H.-S., Ficarro, S.B., Card, J.D., Shim, H., Miduturu, C.V., Simeonov, A., Shen, M., Marto, J.A., Dhe-Paganon, S., Hall, M.D., Cantley, L.C., Gray, N.S., 2020. Discovery and structure–activity relationship study of (Z)-5-methylenethiazolidin-4-one derivatives as potent and selective pan-phosphatidylinositol 5-phosphate 4-kinase inhibitors. J Med Chem, 63, 4880–4895.

Němec, V., Hylsová, M., Maier, L., Flegel, J., Sievers, S., Ziegler, S., Schröder, M., Berger, B.-T., Chaikuad, A., Valčíková, B., Uldrijan, S., Drápela, S., Souček, K., Waldmann, H., Knapp, S., Paruch, K., 2019. Furo[3,2-b]pyridine: A privileged scaffold for highly selective kinase inhibitors and effective modulators of the hedgehog pathway. Angew Chem Int Ed, 58, 1062–1066.

Paquet, T., Le Manach, C., Cabrera, D.G., Younis, Y., Henrich, P.P., Abraham, T.S., Lee, M.C.S., Basak, R., Ghidelli-Disse, S., Lafuente-Monasterio, M.J., Bantscheff, M., Ruecker, A., Blagborough, A.M., Zakutansky, S.E., Zeeman, A.M., White, K.L., Shackleford, D.M., Mannila, J., Morizzi, J., Scheurer, C., Angulo-Barturen, I., Martínez, M.S., Ferrer, S., Sanz, L.M., Gamo, F.J., Reader, J., Botha, M., Dechering, K.J., Sauerwein, R.W., Tungtaeng, A., Vanachayangkul, P., Lim, C.S., Burrows, J., Witty, M.J., Marsh, K.C., Bodenreider, C., Rochford, R., Solapure, S.M., Jiménez-Díaz, M.B., Wittlin, S., Charman, S.A., Donini, C., Campo, B., Birkholtz, L.M., Hanson, K.K., Drewes, G., Kocken, C.H.M., Delves, M.J., Leroy, D., Fidock, D.A., Waterson, D., Street, L.J., Chibale, K., 2017. Antimalarial efficacy of MMV390048, an inhibitor of Plasmodium phosphatidylinositol 4-kinase. Sci Transl Med, 9, eaad9735.

Poli, A., Abdul-Hamid, S., Zaurito, A.E., Campagnoli, F., Bevilacqua, V., Sheth, B., Fiume, R., Pagani, M., Abrignani, S., Divecha, N., 2021. PIP4Ks impact on PI3k, FOXP3, and UHRF1 signaling and modulate human regulatory T cell proliferation and immunosuppressive activity. Proc Natl Acad Sci U S A, 118, e2010053118.

Poli, A., Zaurito, A.E., Abdul-Hamid, S., Fiume, R., Faenza, I., Divecha, N., 2019. Phosphatidylinositol 5 phosphate (PI5P): From behind the scenes to the front (nuclear) stage. Int J Mol Sci, 20, 2080.

Potjewyd, F.M., Marquez, A.B., Chaikuad, A., Howell, S., Dunn, A.S., Beltran, A.A., Smith, J.L., Drewry, D.H., Beltran, A.S., Axtman, A.D., 2022. Modulation of tau tubulin kinases (TTBK1 and TTBK2) impacts ciliogenesis. bioRxiv, 2022.2005.2006.490937.

Preynat-Seauve, O., Nguyen, E.B.-V., Westermaier, Y., Héritier, M., Tardy, S., Cambet, Y., Feyeux, M., Caillon, A., Scapozza, L., Krause, K.-H., 2021. Novel mechanism for an old drug: Phenazopyridine is a kinase inhibitor affecting autophagy and cellular differentiation. Front Pharmacol, 12, 664608.

Ramsden, N., Perrin, J., Ren, Z., Lee, B.D., Zinn, N., Dawson, V.L., Tam, D., Bova, M., Lang, M., Drewes, G., Bantscheff, M., Bard, F., Dawson, T.M., Hopf, C., 2011. Chemoproteomics-based design of potent LRRK2-selective lead compounds that attenuate Parkinson’s disease-related toxicity in human neurons. ACS Chem Biol, 6, 1021–1028.

Rao, V.D., Misra, S., Boronenkov, I.V., Anderson, R.A., Hurley, J.H., 1998. Structure of type IIbeta phosphatidylinositol phosphate kinase: A protein kinase fold flattened for interfacial phosphorylation. Cell, 94, 829–839.

Raychaudhuri, S., Remmers, E.F., Lee, A.T., Hackett, R., Guiducci, C., Burtt, N.P., Gianniny, L., Korman, B.D., Padyukov, L., Kurreeman, F.A., Chang, M., Catanese, J.J., Ding, B., Wong, S., van der Helm-van Mil, A.H., Neale, B.M., Coblyn, J., Cui, J., Tak, P.P., Wolbink, G.J., Crusius, J.B., van der Horst-Bruinsma, I.E., Criswell, L.A., Amos, C.I., Seldin, M.F., Kastner, D.L., Ardlie, K.G., Alfredsson, L., Costenbader, K.H., Altshuler, D., Huizinga, T.W., Shadick, N.A., Weinblatt, M.E., de Vries, N., Worthington, J., Seielstad, M., Toes, R.E., Karlson, E.W., Begovich, A.B., Klareskog, L., Gregersen, P.K., Daly, M.J., Plenge, R.M., 2008. Common variants at CD40 and other loci confer risk of rheumatoid arthritis. Nat Genet, 40, 1216–1223.

Rodgers, G., Austin, C., Anderson, J., Pawlyk, A., Colvis, C., Margolis, R., Baker, J., 2018. Glimmers in illuminating the druggable genome. Nat Rev Drug Discov, 17, 301–302.

Sharma, G., Guardia, C.M., Roy, A., Vassilev, A., Saric, A., Griner, L.N., Marugan, J., Ferrer, M., Bonifacino, J.S., DePamphilis, M.L., 2019. A family of PIKFYVE inhibitors with therapeutic potential against autophagy-dependent cancer cells disrupt multiple events in lysosome homeostasis. Autophagy, 15, 1694–1718.

Shim, H., Wu, C., Ramsamooj, S., Bosch, K.N., Chen, Z., Emerling, B.M., Yun, J., Liu, H., Choo-Wing, R., Yang, Z., Wulf, G.M., Kuchroo, V.K., Cantley, L.C., 2016. Deletion of the gene Pip4k2c, a novel phosphatidylinositol kinase, results in hyperactivation of the immune system. Proc Natl Acad Sci U S A, 113, 7596–7601.

Sivakumaren, S.C., Shim, H., Zhang, T., Ferguson, F.M., Lundquist, M.R., Browne, C.M., Seo, H.S., Paddock, M.N., Manz, T.D., Jiang, B., Hao, M.F., Krishnan, P., Wang, D.G., Yang, T.J., Kwiatkowski, N.P., Ficarro, S.B., Cunningham, J.M., Marto, J.A., Dhe-Paganon, S., Cantley, L.C., Gray, N.S., 2020. Targeting the PI5P4K lipid kinase family in cancer using covalent inhibitors. Cell Chem Biol, 27, 525–537.e526.

Sun, Y., Thapa, N., Hedman, A.C., Anderson, R.A., 2013. Phosphatidylinositol 4,5-bisphosphate: Targeted production and signaling. Bioessays, 35, 513–522.

Wang, D.G., Paddock, M.N., Lundquist, M.R., Sun, J.Y., Mashadova, O., Amadiume, S., Bumpus, T.W., Hodakoski, C., Hopkins, B.D., Fine, M., Hill, A., Yang, T.J., Baskin, J.M., Dow, L.E., Cantley, L.C., 2019. PIP4Ks suppress insulin signaling through a catalytic-independent mechanism. Cell Rep, 27, 1991–2001.e1995.

Wells, C., Couñago, R.M., Limas, J.C., Almeida, T.L., Cook, J.G., Drewry, D.H., Elkins, J.M., Gileadi, O., Kapadia, N.R., Lorente-Macias, A., Pickett, J.E., Riemen, A., Ruela-de-Sousa, R.R., Willson, T.M., Zhang, C., Zuercher, W.J., Zutshi, R., Axtman, A.D., 2019. SGC-AAK1-1: A chemical probe targeting AAK1 and BMP2K. ACS Med Chem Lett, 11, 340–345.

Wells, C.I., Drewry, D.H., Pickett, J.E., Tjaden, A., Krämer, A., Müller, S., Gyenis, L., Menyhart, D., Litchfield, D.W., Knapp, S., Axtman, A.D., 2021. Development of a potent and selective chemical probe for the pleiotropic kinase CK2. Cell Chem Biol, 28, 546–558.e510.

Wortmann, L., Bräuer, N., Holton, S.J., Irlbacher, H., Weiske, J., Lechner, C., Meier, R., Karén, J., Siöberg, C.B., Pütter, V., Christ, C.D., ter Laak, A., Lienau, P., Lesche, R., Nicke, B., Cheung, S.-H., Bauser, M., Haegebarth, A., von Nussbaum, F., Mumberg, D., Lemos, C., 2021. Discovery and characterization of the potent and highly selective 1,7-naphthyridine-based inhibitors BAY-091 and BAY-297 of the kinase PIP4K2A. J Med Chem, 64, 15883–15911.

Wright, B.D., Loo, L., Street, S.E., Ma, A., Taylor-Blake, B., Stashko, M.A., Jin, J., Janzen, W.P., Frye, S.V., Zylka, M.J., 2014. The lipid kinase PIP5K1C regulates pain signaling and sensitization. Neuron, 82, 836–847.

Zhang, Y., Wang, H., Chen, T., Wang, H., Liang, X., Zhang, Y., Duan, J., Qian, S., Qiao, K., Zhang, L., Liu, Y., Wang, J., 2021. C24-ceramide drives gallbladder cancer progression through directly targeting phosphatidylinositol 5-phosphate 4-kinase type-2 gamma to facilitate mammalian target of rapamycin signaling activation. Hepatology, 73, 692–712.

