## Supplementary material for "Identification of a chemical probe for lipid kinase phosphatidylinositol-5-phosphate 4-kinase gamma (PI5P4Kγ)": Combined Supplemental Data

### Table of Contents

Figure S1 S2
Figure S2 S2
Figure S3 S3
Figure S4 S3
Purity tracers and spectra S4–S9

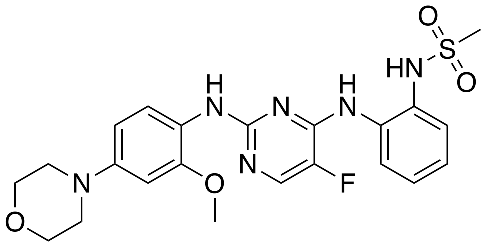

**CZC-25146**

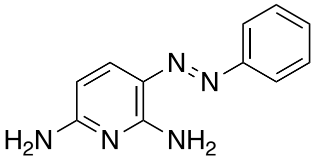

**Phenazopyridine**

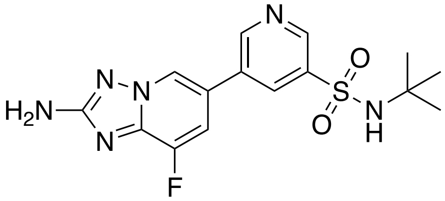

**CZC-24832**

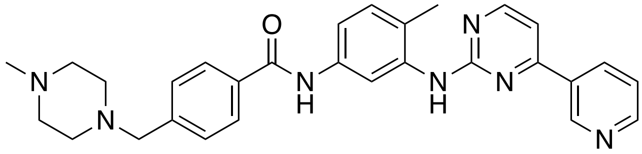

**Gleevec**

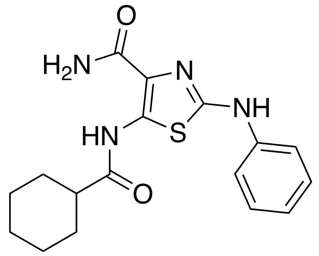

**UNC3230**

**Fig. S1.** Compounds for which PI5P4Kγ is a reported off-target.

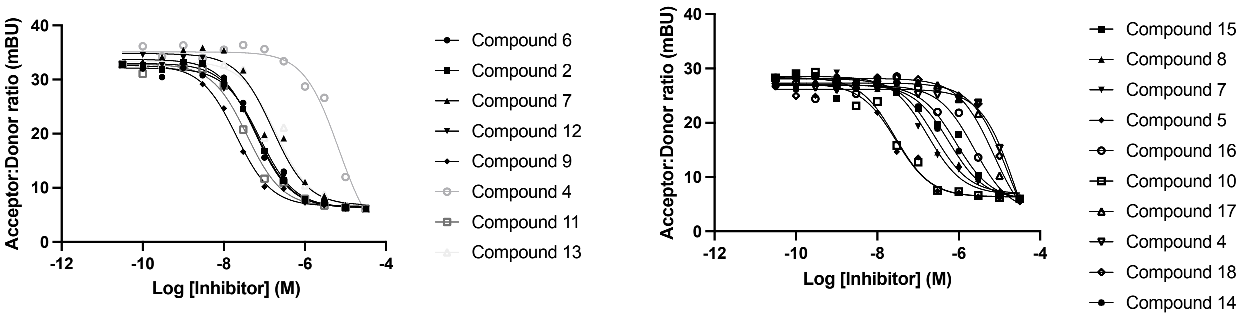


**Fig. S2.** PI5P4Kγ NanoBRET curves for all compounds included.

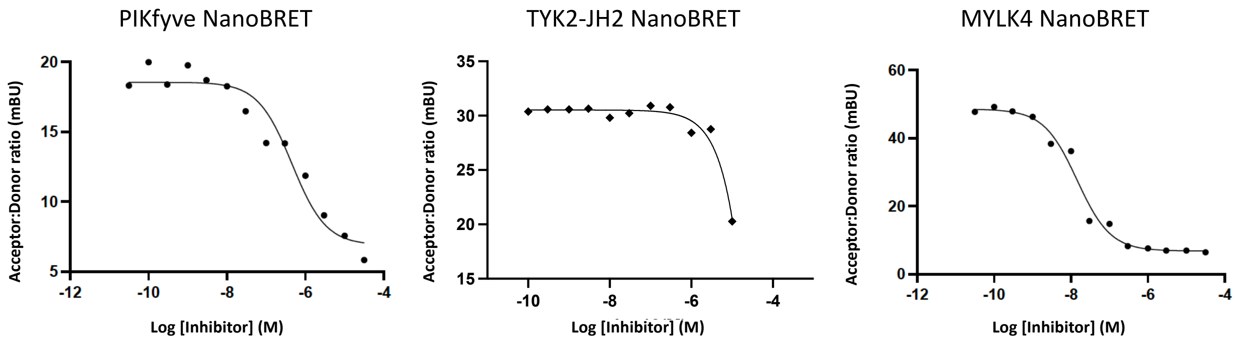


**Fig. S3.** NanoBRET assay curves for off-target kinases when treated with compound **2** in dose-response.

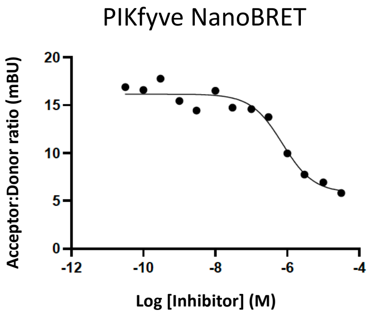


**Fig. S4.** NanoBRET assay curve for PIKfyve when treated with compound **4** in dose-response.

*11-(3,6-dihydro-2H-pyran-4-yl)-5,6,7,8-tetrahydropyrimido[4',5':3,4]cyclohepta[1,2-b]indol-2-amine (**2**)*

**[1]H NMR**

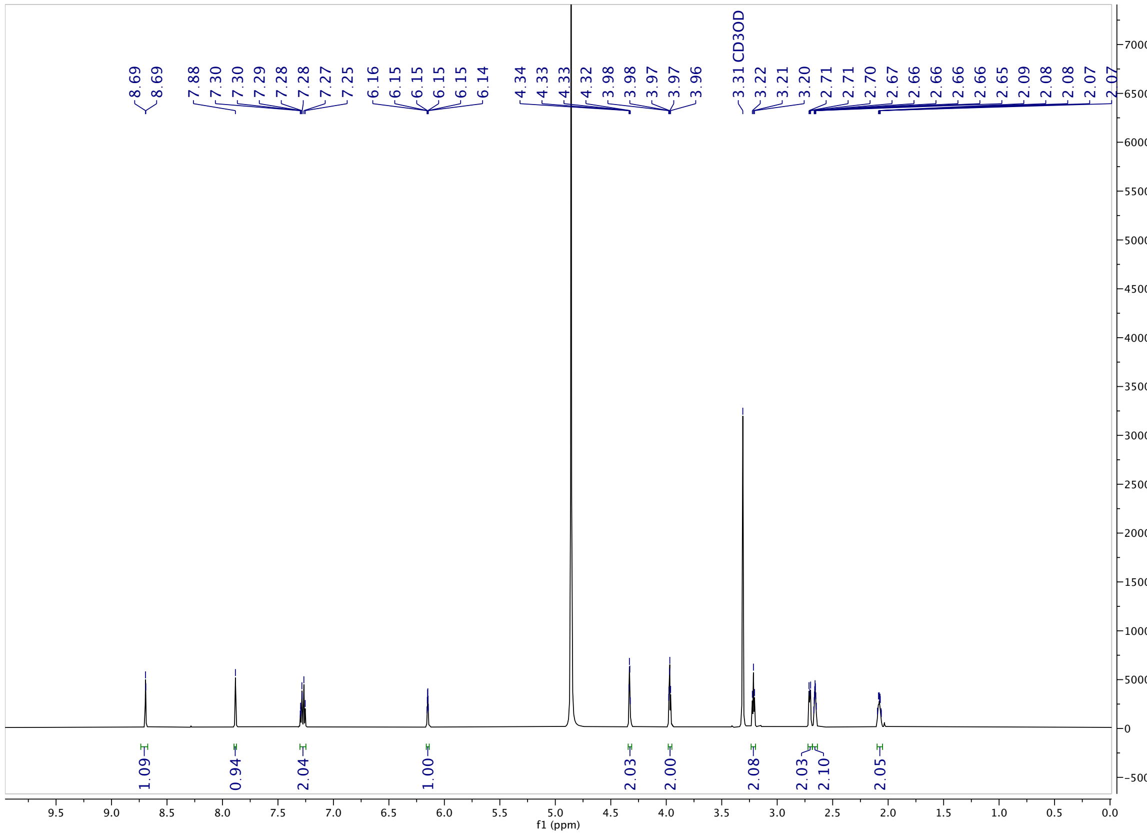

*11-(3,6-dihydro-2H-pyran-4-yl)-5,6,7,8-tetrahydropyrimido[4',5':3,4]cyclohepta[1,2-b]indol-2-amine (**2**)*

**$^{13}$C NMR**

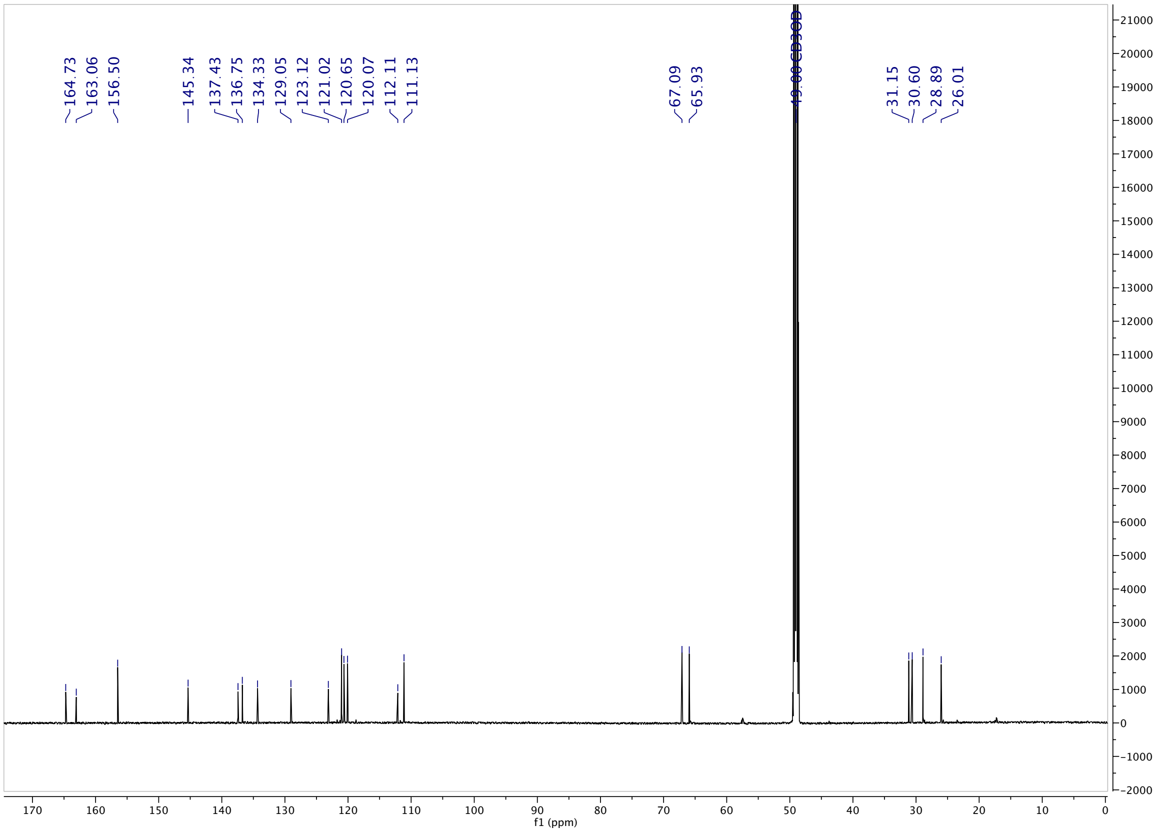

*11-(3,6-dihydro-2H-pyran-4-yl)-5,6,7,8-tetrahydropyrimido[4',5':3,4]cyclohepta[1,2-b]indol-2-amine (**2**)*

### HPLC Purity

MaxPeak: 100.00%
Ret_Time: 0.937 min

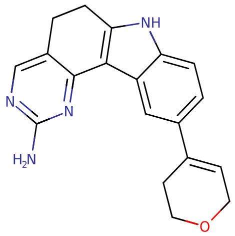


**Mol Wt** 318.37
**Exact Mass** 318.17

| # | Time | Area% |
|---|---|---|
| 1 | 0.937 | 100.00 |

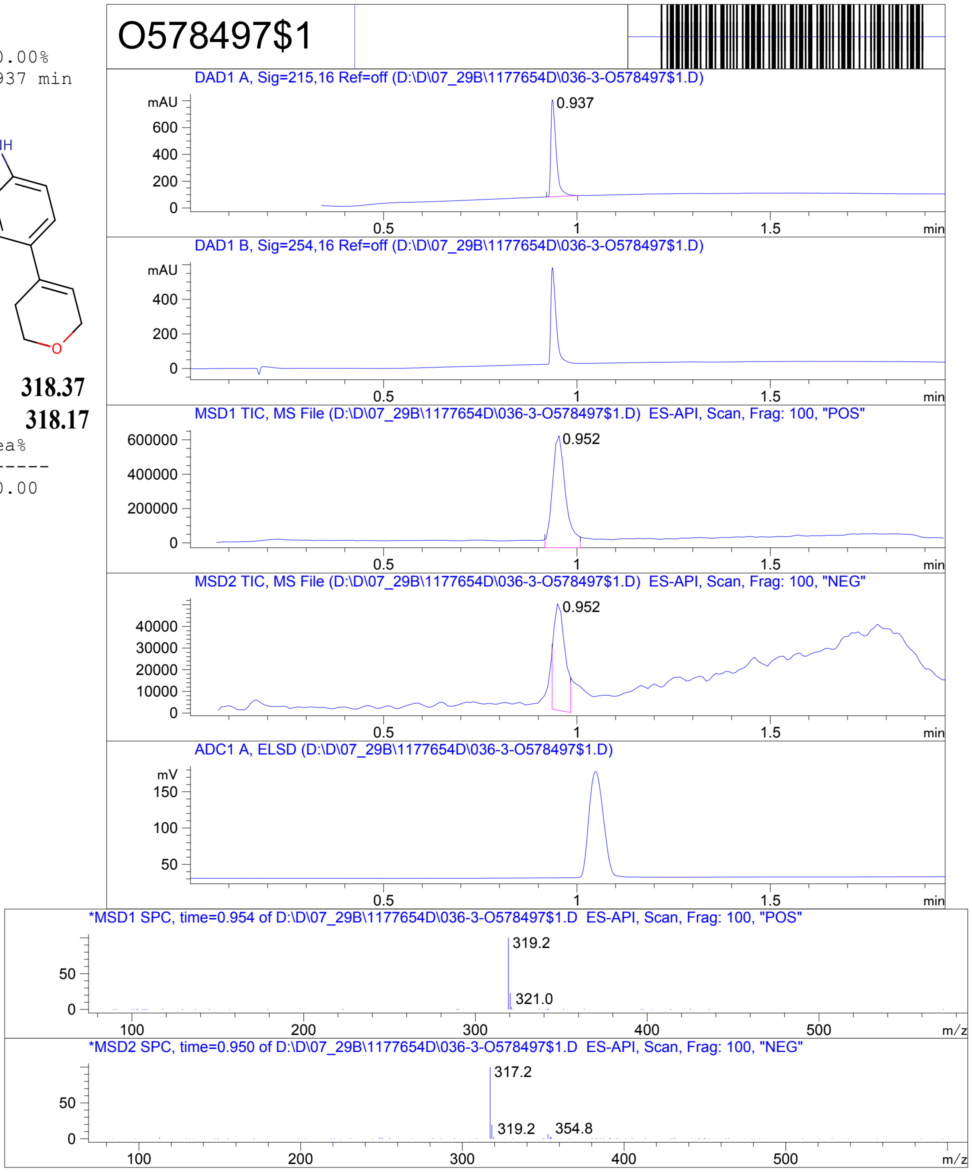


RT 0.952

RT 0.952

*11-(3,6-dihydro-2H-pyran-4-yl)-5,6,7,8-tetrahydropyrimido[4',5':3,4]cyclohepta[1,2-b]indol-2-amine (**2**)*

**HRMS**

| | | | |
|---|---|---|---|
| **Data File** | H1791886.d | **Sample Name** | H1791886 |
| **Sample Type** | Sample | **Position** | P1-A3 |
| **Instrument Name** | Instrument 1 | **User Name** | Denis V.Bylina |
| **Acq Method** | Fast_Gradient_HRMS_pos_Lock_08272019.m | **Acquired Time** | 5/7/2021 6:01:56 PM (UTC+03:00) |
| **IRM Calibration Status** | Success | **DA Method** | Fast_Gradient_HRMS_pos_Lock_08272019.m |
| **Comment** | | | |
| **Sample Group** | | **Info.** | |
| **MFC** | C20H20N4O | **Stream Name** | LC 1 |
| **Acquisition Time (Local)** | 5/7/2021 6:01:56 PM (UTC+03:00) | **Acquisition SW Version** | 6200 series TOF/6500 series Q-TOF B.08.00 (B8058.0) |
| **TOF Driver Version** | 8.00.00 | **TOF Firmware Version** | 8.643 |
| **Tune Mass Range Max.** | 1700 | | |

**Compound Table**

| Label | Tgt Score | Mass Error (ppm) | Tgt Formula | Obs. RT | Ref. Mass | Obs. Mass |
|---|---|---|---|---|---|---|
| Cpd 1: C20 H20 N4 O; 2.250 | 79.22 | 1.1 | C20 H20 N4 O | 2.25 | 332.1637 | 332.1641 |

| Obs. *m/z* | Obs. RT | Obs. Mass | Tgt Formula | Tgt Mass | Tgt Mass Error (ppm) | RT Diff. | Find Cpds Algorithm |
|---|---|---|---|---|---|---|---|
| 333.1709 | 2.25 | 332.1641 | C20 H20 N4 O | 332.1637 | 1.1 | Find By Formula | |

**Compound Chromatograms**

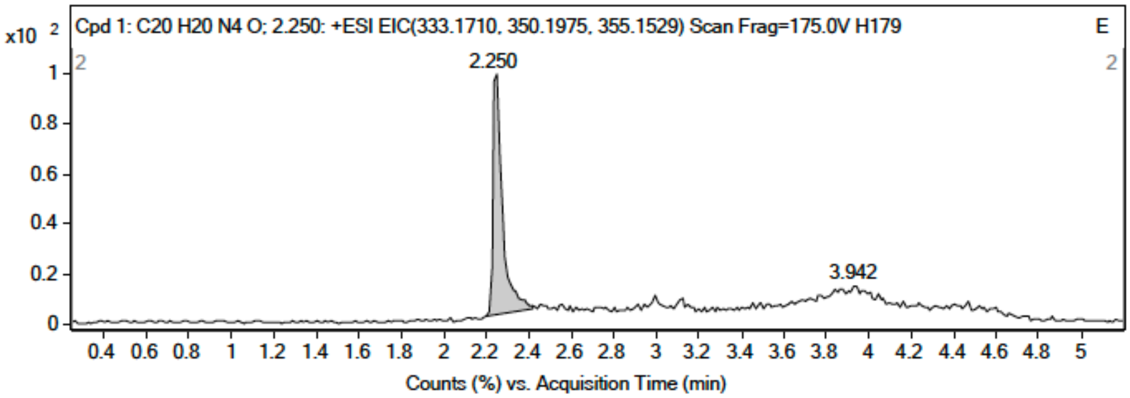

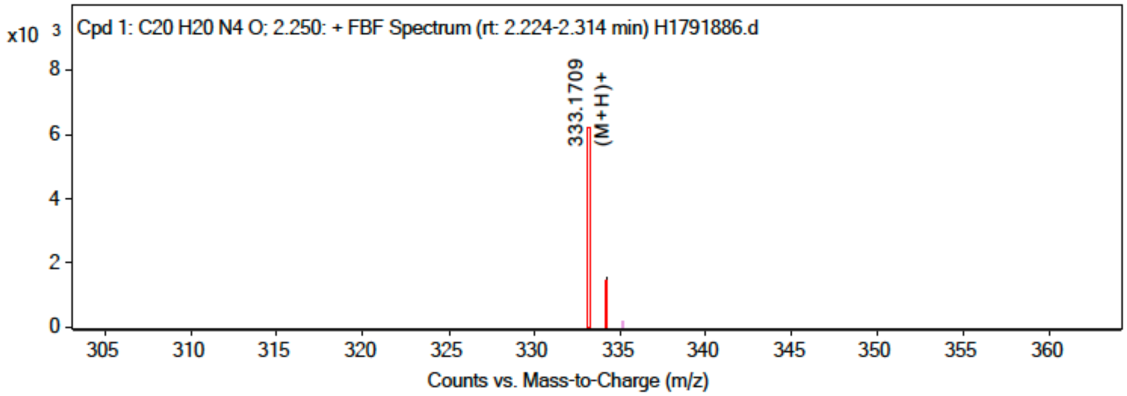


**MS Spectrum Peak List**

| Obs. *m/z* | Charge | Abund | Ion/Isotope |
|---|---|---|---|
| 333.1709 | 1 | 6136.24 | (M+H)+ |
| 334.1761 | 1 | 1549.18 | (M+H)+ |

MS Zoomed Spectrum

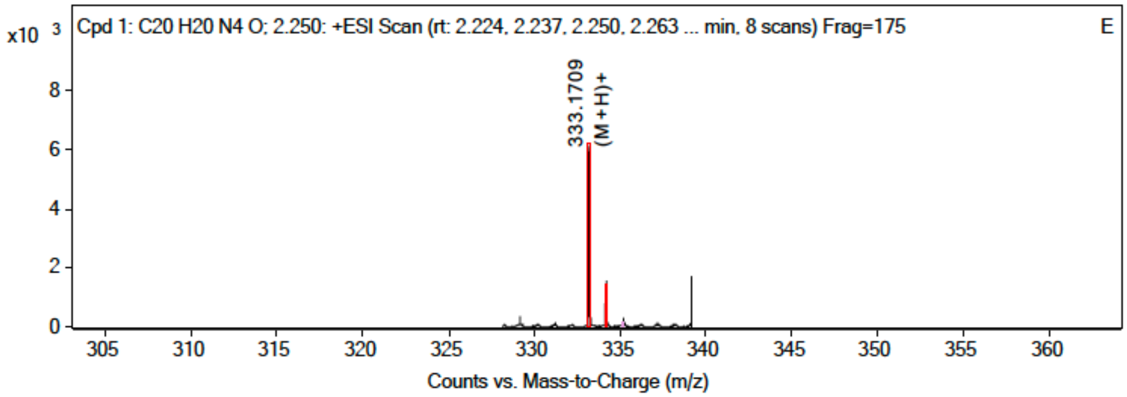


**MS Spectrum Peak List**

| Obs. *m/z* | Charge | Abund | Ion/Isotope | Tgt Mass Error (ppm) |
|---|---|---|---|---|
| 333.1709 | 1 | 6136.24 | (M+H)+ | 0.2 |
| 333.1709 | | 6136.24 | | |
| 334.1761 | 1 | 1549.18 | (M+H)+ | -6.2 |

*12-(cyclopropylethynyl)-6,7,8,9-tetrahydro-5H-pyrimido[4',5':3,4]cycloocta[1,2-b]indol-2-amine (**4**)*

### $^{1}$H NMR

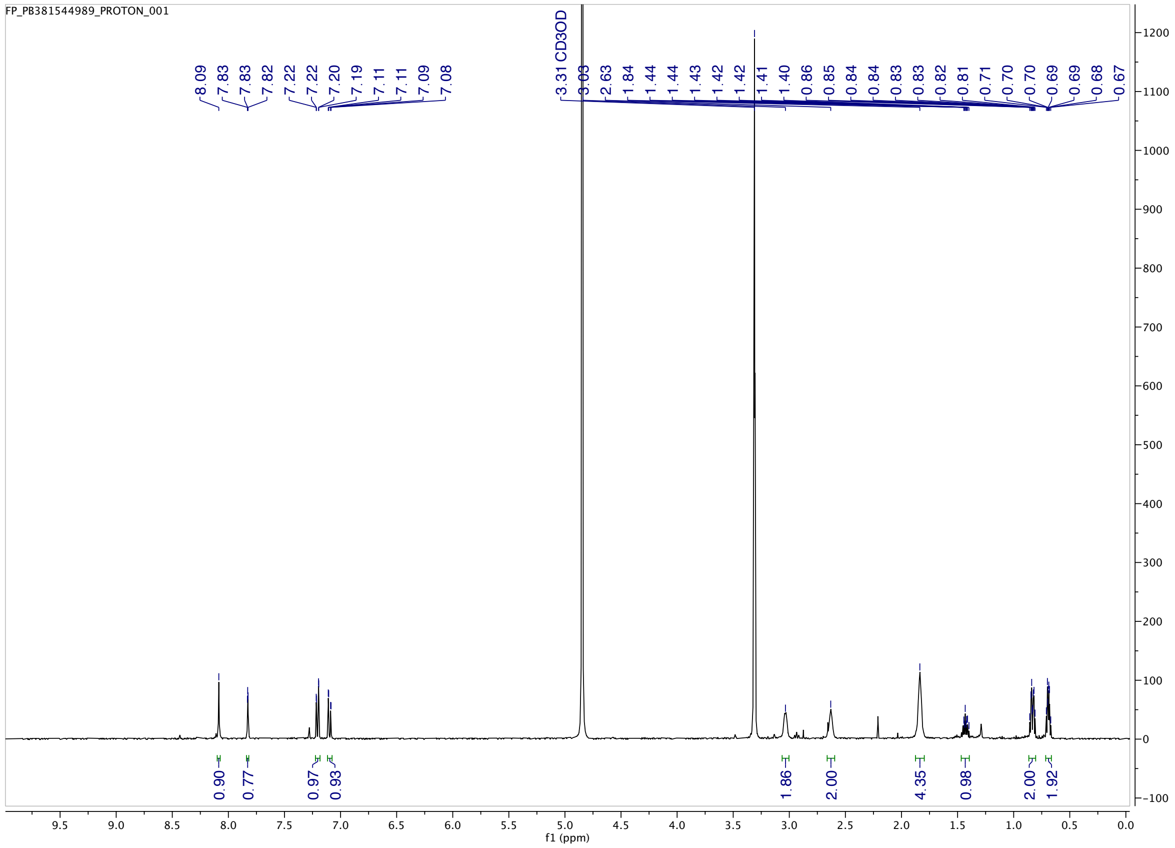
